# Improving quantitative structure models of trees inspired by pipe and metabolic scaling theory

**DOI:** 10.1101/2022.10.31.514601

**Authors:** Jan Hackenberg, Jean-Daniel Bontemps

## Abstract

**Purpose:** We invent in this manuscript new tree parameters which can be derived from a single QSM. QSMs are topological ordered cylinder models of trees which describe the branching structure up to the tips. All new invented parameters have in common, that their defining point of view looks from the direction of the tips and not from the root along the tree.

**Methods:** We use new allometric power functions to predict the radius from the invented parameters. Then we improve the radii of the QMSs’ cylinders utilizing those functions.

**Results:** For validation we use QSMs produced from an open point cloud data set of tree clouds with SimpleForest software. We compare the QSM volume against the harvested reference data for 65felled trees. We also found QSM data of TreeQSM, a competitive and broadly accepted QSM modeling tool. Our RMSE was less than 40 % of the TreeQSM RMSE. For other error measures, the r^2^_adj._ and the `CCC`, the relative improvement looked even better with reaching only 27 % and 21 % of the TreeQSM errors respectively.

In a second validation part we show a way to run numerical tests against the West Brown Enquist (WBE) model. Expected power coefficients have been published for various allometric relations and we compare them to predicted values from QSM data. The deviation from the expected values ranges here from 8 % underestimation to 32 % overestimation.

**Conclusions:** `Quality -` With the invention of the QSM radius filter technique we improve tree volume prediction capabilities utilizing QSMs.

`Quantity -` More data can be collected with QSMs than with traditional methods.

## 1 Introduction

The scope of this manuscript is the invention of new tree describing parameters which have better correlation with radius/diameter measurements at any point of a tree compared to traditional tree describing parameters. We run automated methods on 3d point cloud data using those parameters to build an efficient filter for volume predictions. We use here terrestrial laser scan (`TLS`) data. From `TLS` clouds we build quantitative structure models (`QSM`) - topological ordered cylinder models describing the branching structure of trees [1–3]. On those `QSM`s we explore new tree modeling techniques. We wrap new methods into a context of traditional forest ecological theories.

Before we use the proposed protocol we want first look back in time to get an insight how efficient the digital method is compared to traditional analysis. Many ways to predict the above ground volume of trees with mathematical concepts have been performed over the last centuries. Already 500 years ago Leonardo da Vinci stated that the sum of cross sectional areas at any height equals the cross sectional area of the stem [4]. From this hypothesis a `Volume` prediction model utilizing the Diameter at Breast Height (`DBH`) and the Height (`H`) follows, see equation 1.

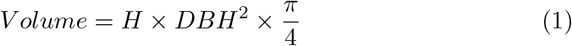

While we know this equation overestimates the `Volume` it is a quite fundamental ecological observation based statistical tree model because the cylinder formula is a really simple geometric form. Additionally it relies only on two easy to obtain field parameters.

Also in da Vinci’s work firstly the comparison of a tree’s branching system with a river network was performed. An important ordering attribute describing river networks is the number of sources feeding that part of the river. We will discuss in this manuscript expressing branch orders in a reverse and recursive way. In fact one of our tree parameters discussed is the pendant to that river network measure.

An important explanation for the overestimation caused by equation 1 is the lack of a component which describes the tapering of stem or branch segments. All segments, that is tree parts between two neighboring branch junctions, have the tendency to decrease their diameter if we look from closest to the root to furthest from the root along the segment.

Due to data limitations often only partial volume predictor functions exist in a published form. Branch volume is often ignored and we can for example only express the tree’s stem volume with formulas as we can see in equation 2. The tree species specific form factor *f* describes the taper of the stem and is published for various species [5].

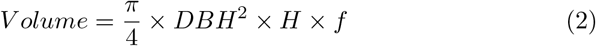

We can find those and other similar tree species specific equations in online data bases [6, 7]. Nevertheless, utilizing those equations comes with a large uncertainty about the prediction accuracy, especially if regional growth pattern are accounted to. With traditional methods the `Volume` could only be destructively collected with high field costs. To retrieve a `Volume` estimate for a single tree the tree had to be felled, sawed into segments, dried and weighted. The total above ground biomass (`AGB`) is then multiplied with the density *ρ*.

### 1.1 Allometric Theory

An interesting `Volume` predictor function for trees is the commonly used power function in equation 3.

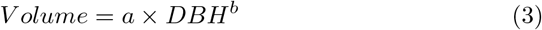

The function is interesting in the sense that it describes the total `Volume` of above ground wood components including branching structure.

Using power functions to model size relations is not limited to trees. The theory of allometric scaling relying on exactly that base function is a general biological theory, see equation 4 for the generalization.

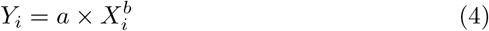

A century ago Huxley analysed size relations on the data of shore-crab bodies and realized he could linearize the problem by a simple log transform on predictor and predicted, as we can see in equation 5 [8]. The error *ϵ*_*i*_ follows a normal distribution with mean 0 and standard deviation *σ*^2^. Huxley assumed an homoscedastic error constant over the full range of log transformed data, see equation 5.

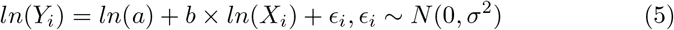

After the linear fit the back transformation to the power function shown in equation 6 was performed by Huxley.

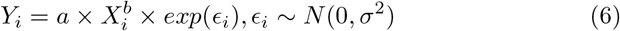

However, the variation in the presented model now is a multiplier for the mean function, which means that the residuals of *Y* will increase as *X* increases. In other words, the distribution for the response variable in the model will be heteroscedastic.

The hypothesis that a part of an organism will generally grow in proportion to some constant power of the size of the whole organism was applied in other works in the following years [9].

One limitation of the allometric scaling analysis protocol published by Huxley [8] was that the protocol required data which had a linear pattern in the logarithmic domain. Various data sets however did not fullfill this precondition but rather had curvilinear form in the logarithmic domain. For a while such data could not be modeled with the allometric scaling theory.

Fifty years after Huxley’s early work science started to apply non linear regression on allometric scaling data which was not logarithmic transformed [10, 11].

Their non linear solver relied on a homoscedastic error, see equation 7.

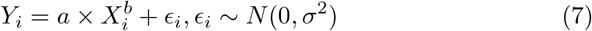

This approach was received critically at the biology audience in the first decade [12–16]. The reasoning of the conservative reception was about the fact that the size of residuals of response *Y*_*i*_ typically increased substantially with the value for *X*_*i*_. So a model relying on a heteroscedasticity domain error was preferred.

We get more review details on allometric scaling, its different development over the last century [17].

After giving an overview in the first part of the manuscript Packard then presents a well designed protocol to analyze 22 different statistical functions relying on the base equation 3. The used data consists of ∼90 individuals of the bivalve mollusk *Thracia phaseolina* [18].

All functions do rely on advancements done to Huxley’s work in the last century. Differences in the models are caused by different assumptions on the error distribution. In a first step Packard considers applying the original protocol, e.g. log transform the data and then perform a linear fit [8] relying on equation 5.

information criterion (AIC) [19, 20] received by applying both a first order polynomial and a second order polynomial fit he considers the tested data as inappropriate for this method as he concludes the better performing second order polynom reveals a curvilinear log scaled domain. So the non linear approach to fit a function is used from that decision on.

We gained now insight into allometric scaling basic concepts but want to turn our focus to trees again. In fact the generic allometric scaling concept can be applied to trees as well.

### 1.2 Pipe Model Theory

Before the invention of the allometric scaling protocol Pressler proposed a new tree modeling theory for deciduous temperate species referred to as Pressler’s law [21]. The law states that the summed up cross sectional areas of pipes formed in the latest growing season correlates to the annual diameter increment of the stem. This hypothesis was concluded from the fact that new leaves have to be fed with nutrients via new pipes building this diameter increment.

Combined with allometric scaling this theory was then even further optimized by accounting to inactive pipes as well in the Pipe Model Theory (PMT) [22, 23]. Due to thinning and self thinning pipes become inactive and build up the heartwood. The inactive pipes are located in the center of the stem and branching units. The active pipes which feed the leaves with nutrients build up the outer perimeter of the wooden structure. We can call this outer part sapwood.

Recently a well written summary about the PMT and related tree describing theories was published [24]. The PMT fulfills some conceptual properties:

1. Sapwood area and leaf area/mass are proportional. Here Pressler’s law is echoed.
2. The area-preserving rule assumes that the conductive sapwood area of a stem at a given height is equal to the cumulative basal area of its daughter axes above that height. We can see here a reflection of the da Vinci rule.
3. The operating time of pipes is limited to the time frame of a year, the nutrient transporting pipes are renewed every year.

### 1.3 West Brown and Enquist model

Various biological theories exist about the allometric exponential parameter *b* in equation 4. The parameter often takes the form of a quarter-power [25–27].

A specialization of the theory is the West, Brown and Enquist (WBE) model [28–32]. The WBE model is designed to be a generic biological model not limited to trees, but this time the data used to publish the theory was data originated in forest ecology. Namely the data already used fifty years ago in the publication of the PMT [23]. The WBE model states that the annual growth in biomass scales as the 3/4 power of body mass. So the authors here fix the exponential *b* parameter in equation 4 to a constant value. They claim that such constants can be applied to everything, even the packaging of species in communities [33].

The WBE model sees trees as branching networks that [34]:

1. supply the leaves contained in a fixed three-dimensional volume with nutrients
2. the energy required to perform the supply chain is minimized
3. the leaves and tips of the network are self-similar

The WBE model is up to date a really comprehensive model but it has also been contested on theoretical and empirical grounds [35, 36].

### 1.4 Tree data availability for allometric scaling research

We have now learned about a few relevant ecological theories. We discussed the complex data collection protocols which are time consuming and costly. But in the context of digitalization also new measurement devices have been developed and they already have been used successfully in the field of forest ecology. We mentioned LiDAR techniques and their ability to represent the surface of the environment with point clouds.

Static terrestrial LiDAR sensors provide in forest environments data which covers the surface of trees with point clouds. For the processing of Terrestrial Laser Scanning (TLS) data 3d tree reconstruction algorithms have been developed [1, 2, 37–40]. Because of the automation level the human bias is minimized during data collection. Tree surfaces can easily be covered with multiple millions of euclidean points when TLS data are recorded.

Since around a decade various software tools to produce so called Quantitative Structure Models (QSMs) have been published [41]. QSMs consist of thousands and sometimes even ten or hundred thousands cylinders [3]. With the complete branching structure being geometrically captured the `Volume` can be measured. Full automation in the modeling of the same data will lead to same QSMs not affected by the person running the computation. Data collection time is roughly in the time frame of few hours or days when collecting TLS data which covers up to multiple hectares of forest.

In addition to the geometrical description also topological information is stored inside a QSM. The order from the tree root to the branch tips is added to the geometry parameters. Figure 1 shows a real QSM of a large tree’s point cloud with more than 35 thousand cylinders. And in figure 2 we draw an idealized and simplified mathematical QSM. We discuss the topological order represented by the informatics tree structure later to create new relevant tree parameters.

**Fig.1.**
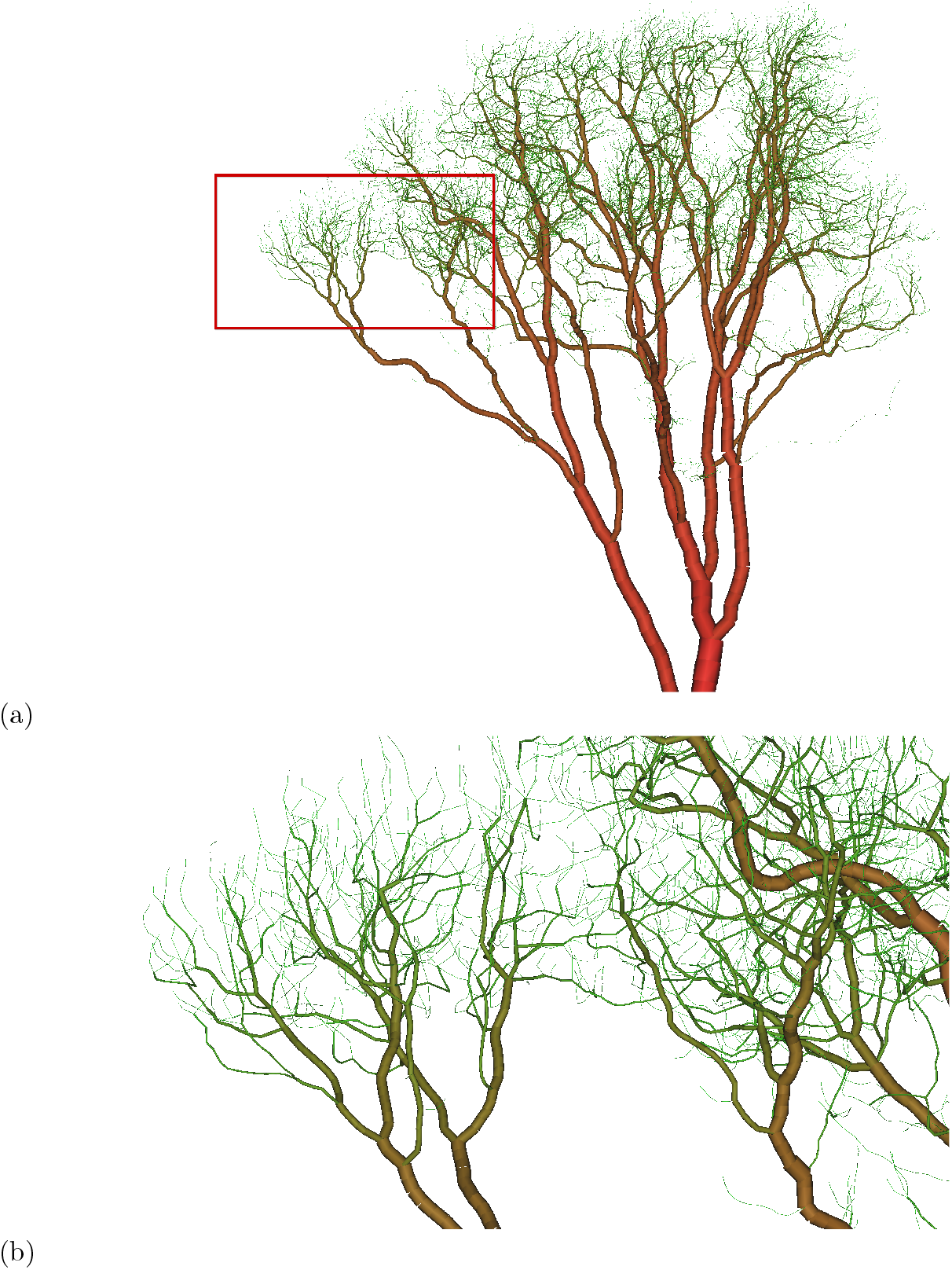
(a) A QSM of an *Acer pseudoplatanus* [47, 48] with a DBH of ∼1.4 *m* containing approximately 25 *m*^3^ of woody biomass. The QSM was modeled with Simple Forest [44]. The tree is located in Wytham Woods, UK. The QSM consists of more than 35 000 cylinders. (b) A zoom of the red bounding boxed area in (a).

**Fig. 2.**
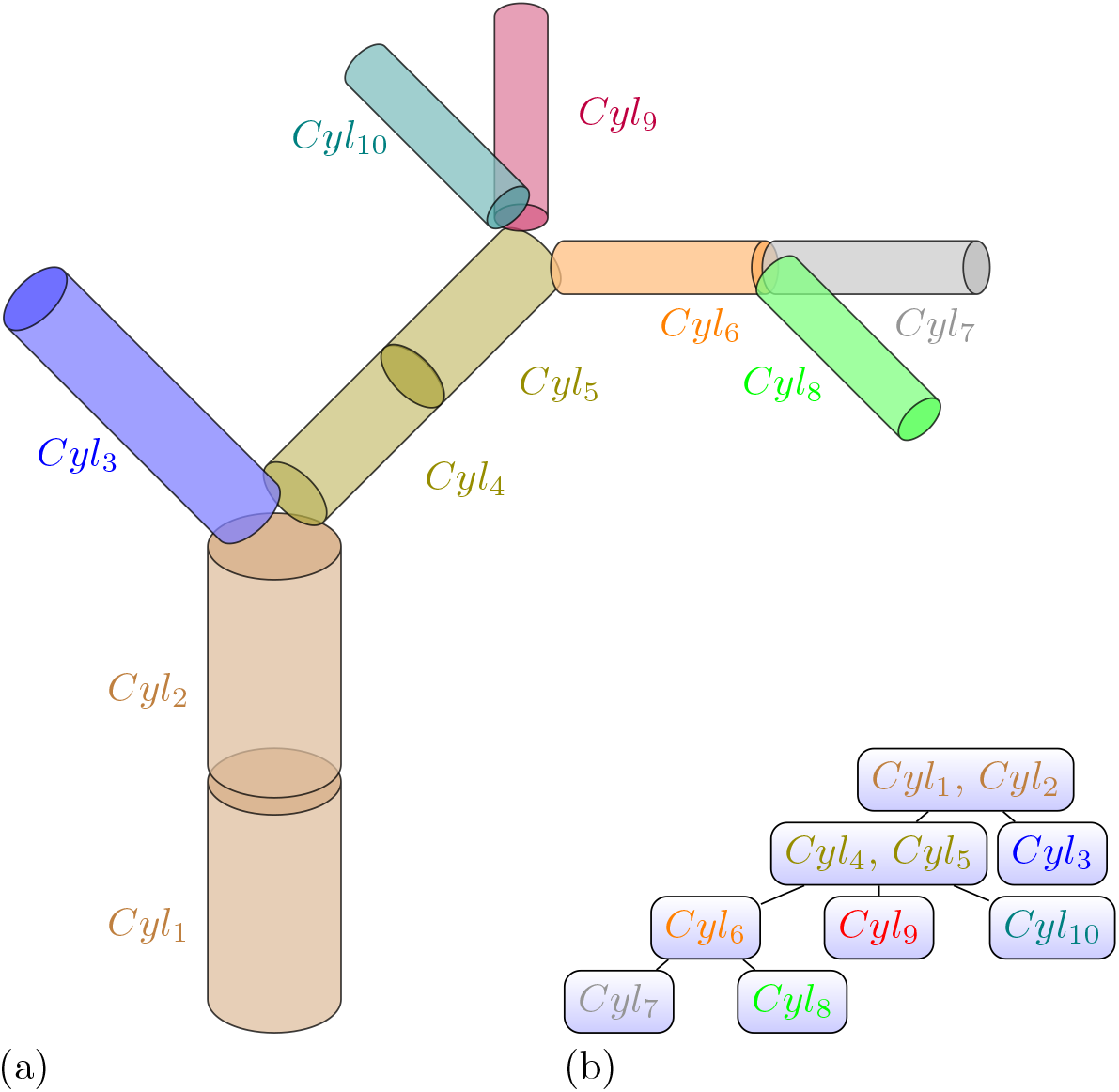
(a) An artificial QSM consisting of ten cylinders Cyl_n_. Each cylinder consists of two 3d points `start` and `end` as well as the `Radius`. The cylinders between two neighboring branching junctions are combined to a list structure called segment. Each segment is colored differently. The cylinder list is always ordered from closest to the root to farthest away from the root. (b) We can see the topological order information from the QSM in (a). The segments are stored in the nodes. The root node is the segment which stores the root cylinder at its begin.

Licensed under copy left are the Computree [42] plugins SimpleTree [2, 43] and SimpleForest [44]. Also TreeQSM [1, 45] and 3dForest [46] are QSM producing tools which can be downloaded with their source code being free.

The free tools have been tested by comparing QSM estimated `Volume` derived from TLS data versus destructive collected `Volume` field measurements [41, 44].

## 2 Methods

First we define new tree describing parameters and then we will build with two of them two functions which can predict the radius of a cylinder by looking at the topological information of the cylinder. We will use both functions in `Radius` filters designed to improve QSM data.

### 2.1 Tree describing parameters

We will get aware of limitations by reading a QSM in the traditional way only. Traditional parameters like the `Branch Order` (`BO`) can be extracted. But we also have the potential to extract new yet untested and undiscussed parameters.

#### 2.1.1 Branch Order

The `BO` is counted from the stem up to the tips.

#### 2.1.2 GrowthVolume

We defined in [43] the `GrowthVolume` of a cylinder recursively.

##### Definition 1

(GrowthVolume) The `GrowthVolume` of a cylinder is the cylinder’s volume plus the `GrowthVolumes` of its children.

In other words the `GrowthVolume` is the sum of the `Volumes` of all cylinders with a child relation to the query cylinder, see figure 3.

**Fig. 3.**
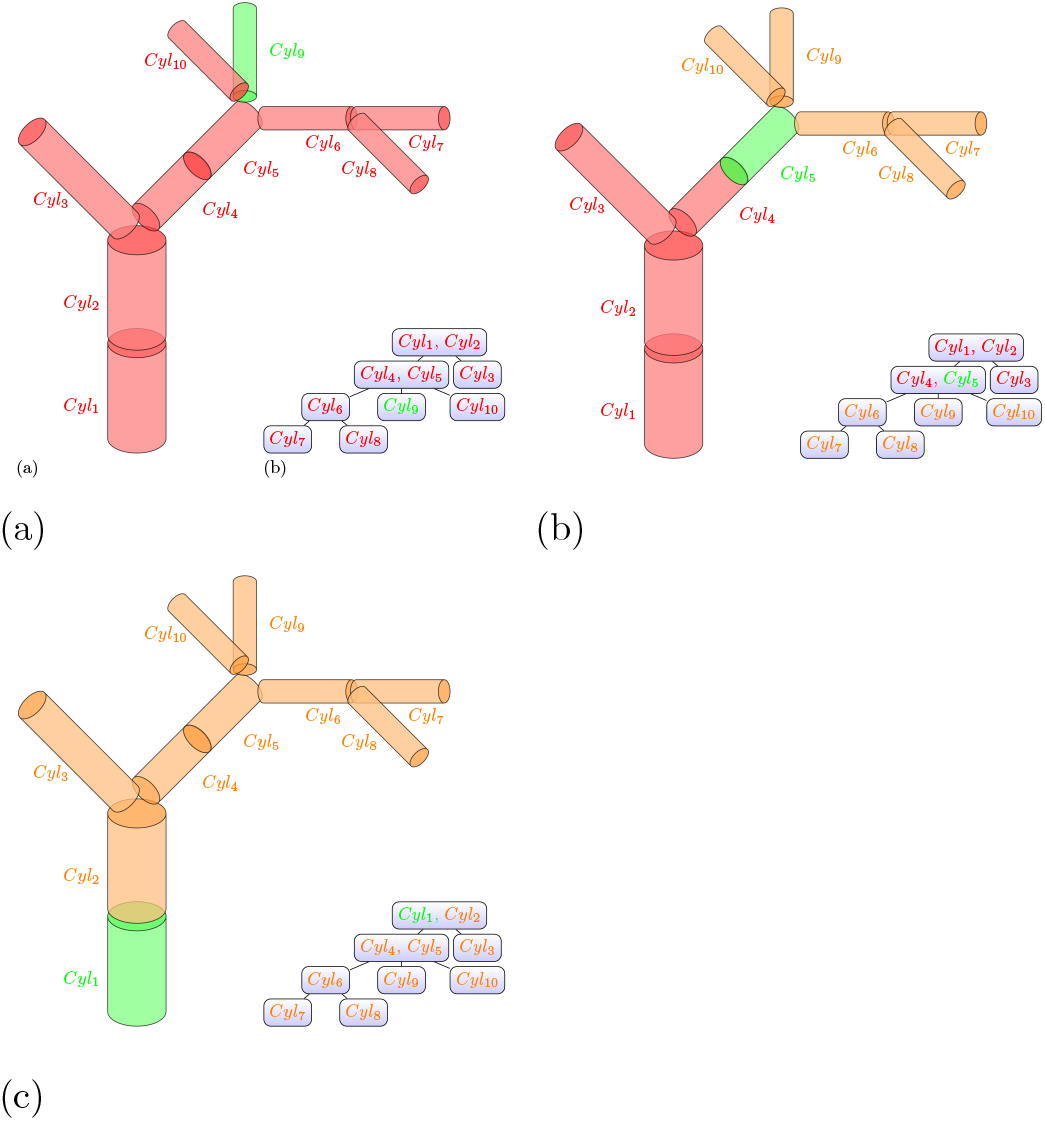
We see a QSM colored in three different colors representing the concept of recursive child relation for a cylinder. In green the query cylinder is highlighted. The query cylinder has a recursive child relation to itself. Orange are all cylinders which have additionally a recursive child relation to that query cylinder. Red are all cylinders colored which have no recursive relation to the query cylinder. Three border cases are depicted: (a) The query cylinder *Cyl*_9_ is a tip node and no other cylinder has a recursive child relation. (b) Here the query cylinder *Cyl*_5_ is located in the center of the QSM. The cylinders fullfilling the the child relation is the sub-branch which would be received when cutting the represented tree at the position between *Cyl*_4_ and *Cyl*_5_. (c) The query cylinder *Cyl*_1_ is the root cylinder of the QSM. The root cylinder’s recursive children are all cylinders of the QSM.

We used the `GrowthVolume` as a predictor for the radius in a radius correction filter [43], see equation 8.

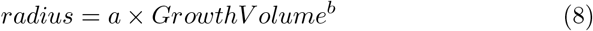

Nevertheless this filter has a design flaw. Overestimated `Radii` of cylinders belonging to the `GrowthVolume` part affect their `Volumes` and therefore also the `GrowthVolume`. Our predictor is affected by the noisy attribute we want to get rid of and we will propose here later a replacement for the `GrowthVolume` not having that design flaw.

#### 2.1.3 GrowthLength

We can substitute the volume with the length in our previous definition to get a recursively defined length.

##### Definition 2

(GrowthLength) The `GrowthLength` of a cylinder is the cylinder’s length plus the `GrowthLength` of its children.

As the `GrowthLength` is not affected by radius prediction errors during cylinder fitting it is a robust ordering predicate for the order of child nodes. With this ordering we can extract the stem by following the left path in the QSM structure depicted in the figure 2.

#### 2.1.4 Reverse Branch Order

To define another meaningful order in the sense of diameter scaling we can read the inverse branch order from a tree.

##### Definition 3

(Reverse Branch Order) The `Reverse Branch Order` (`RBO`) of a cylinder is the maximum depth of the subtree of the segment’s node. If we look at the real tree, the `RBO` denotes the maximal number of branching splits of the sub-branch growing out the segment.

The reverse branch order is depicted in the schematic QSM in figure 4 b).

**Fig. 4.**
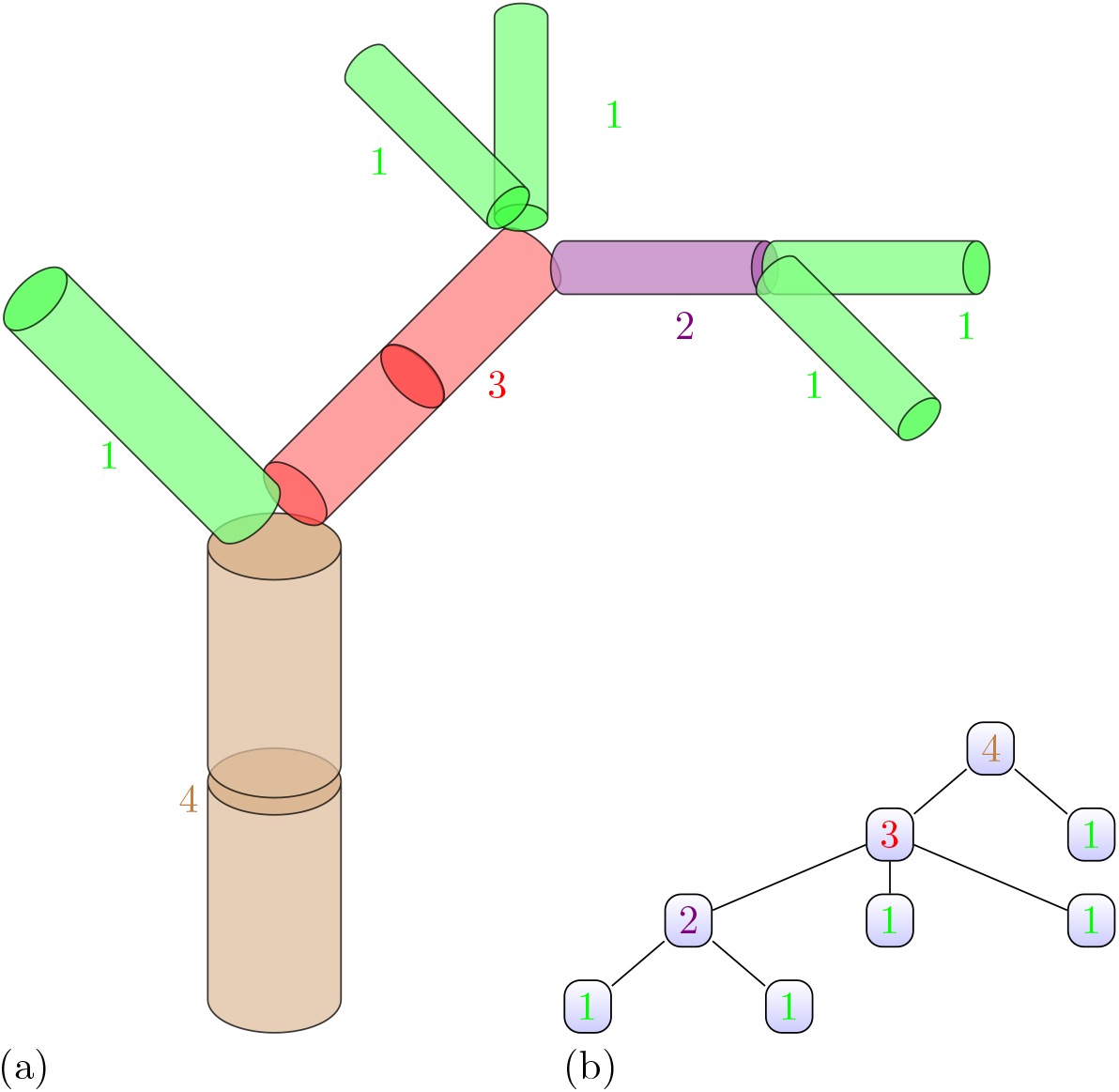
(a) The artificial QSM of figure 2 labeled with the `RBO`. (b) We can see the topological order information from the QSM in (a) and each node has the `RBO` displayed.

#### 2.1.5 Reverse Branch Order Pipe Area

##### Definition 4

(Reverse Branch Order Pipe Area) We define the Reverse Branch Order Pipe Area (`RBOPA`): The WBE model states that the branching architecture is self-similar [49]. In a self-similar branching architecture the `Radius` as well as the cross sectional area of a tip is constant. The `RBOPA` is a tree specific area unit. The expected cross sectional area of each tip of a tree equals `AreaOfTip` *cm*^2^. `AreaOfTip` is later crossed out in our calculus so we do not have to compute it in the following. By following the pipe model theory [23] the `RBOPA` of a cylinder at any location inside the QSM equals the number of supported tips. Supported tip cylinders always have a recursively defined parent relation to the query cylinder.

We can see the `RBOPA` depicted in the schematic QSM visible in figure 5.

**Fig. 5.**
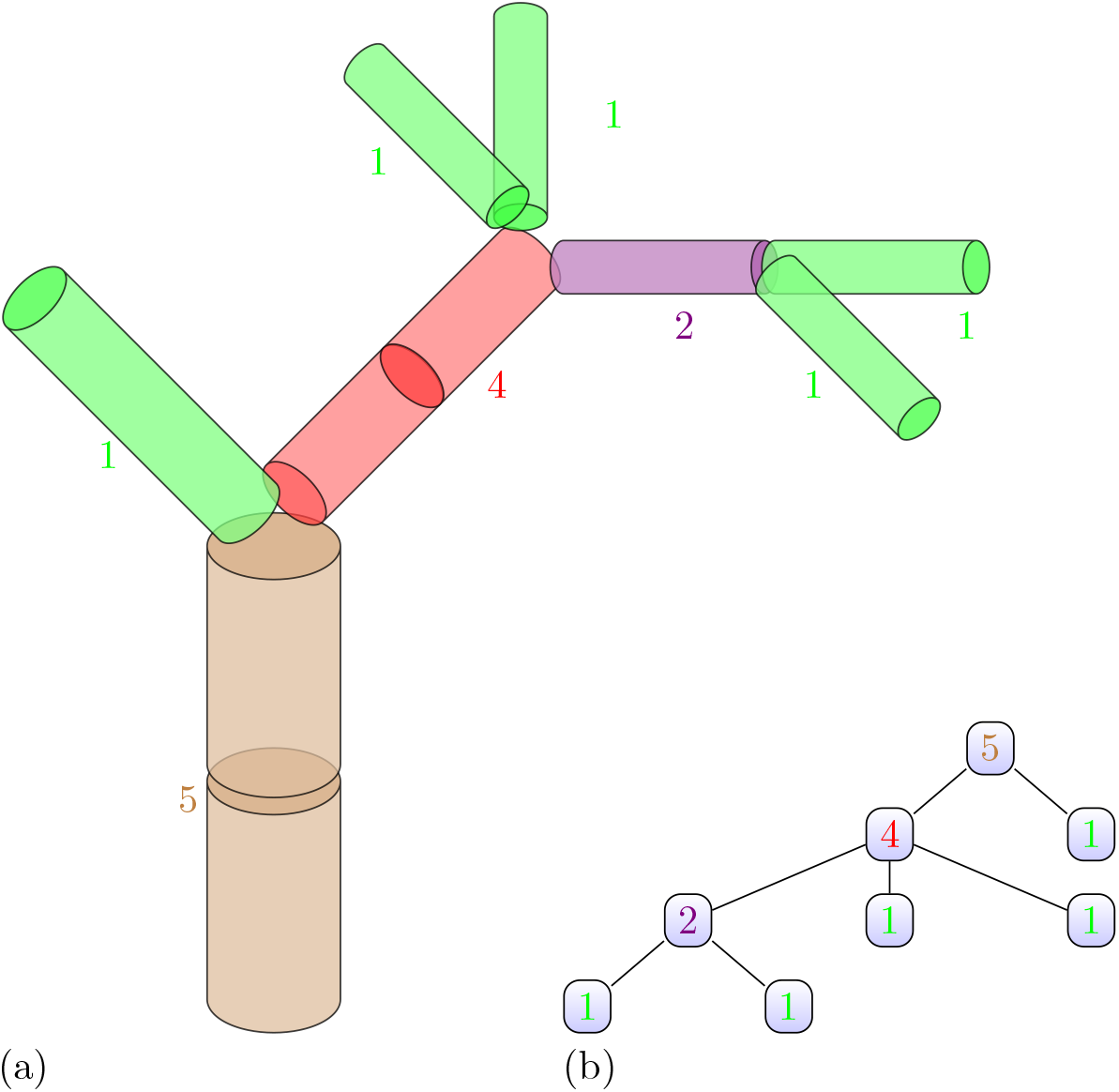
(a) The artificial QSM of figure 1 labeled with the `RBOPA`. (b) We can see the topological order information from the QSM in (a) and each node has the `RBOPA` displayed.

By its definition the `RBOPA` is a proxy for the sapwood area. To derive the true sapwood area measured in metrical units we need to simply multiply `RBOPA`*×* `AreaOfTip`.

#### 2.1.6 Reverse Branch Order Pipe Radius

Earlier works have shown that small cylinders can represent the topology correctly, but their radius tends to be overestimated [50, 51]. By having a radius independent proxy for the cross sectional area we can simply take its square-root to derive a proxy for the radius of a cylinder.

##### Definition 5

(Reverse Branch Order Pipe Radius) We define the Reverse Branch Order Pipe Radius (`RBOPR`) as the square root of the `RBOPA`.

#### 2.1.7 ProxyVolume

As the `RBOPA` is a proxy for the area of active vessels which pass the cylinder we can also model a proxy for the volume occupied by vessels in a cylinder.

##### Definition 6

(ProxyVolume) The `ProxyVolume` of a cylinder is calculated by multiplying the `Length` with the `RBOPA` as shown in equation 9. By definition the `ProxyVolume` is not relying on correctly modeled `Radii`.

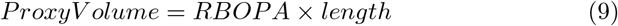

#### 2.1.8 VesselVolume

We had defined the `GrowthVolume` of a cylinder in a recursive manner. We used the `GrowthVolume` in earlier works [43] as a predictor for a corrected radius of each cylinder utilizing a simple power function fit. In similar manner to the `GrowthVolume` we can define the `VesselVolume`, but utilizing instead of cylinders’ `Volumes` their `ProxyVolumes`.

##### Definition 7

(VesselVolume) The `VesselVolume` of a cylinder is its `ProxyVolume` plus the `VesselVolume` of the cylinder’s children.

### 2.2 Comparison of parameters

In branch order one, not only the largest branch segment is included, but also medium sized branches and even epicormic shoots. Therefore the correlation between this branch order and the diameter of segments of a certain branch order is rather low, see figure 6 a).

**Fig. 6.**
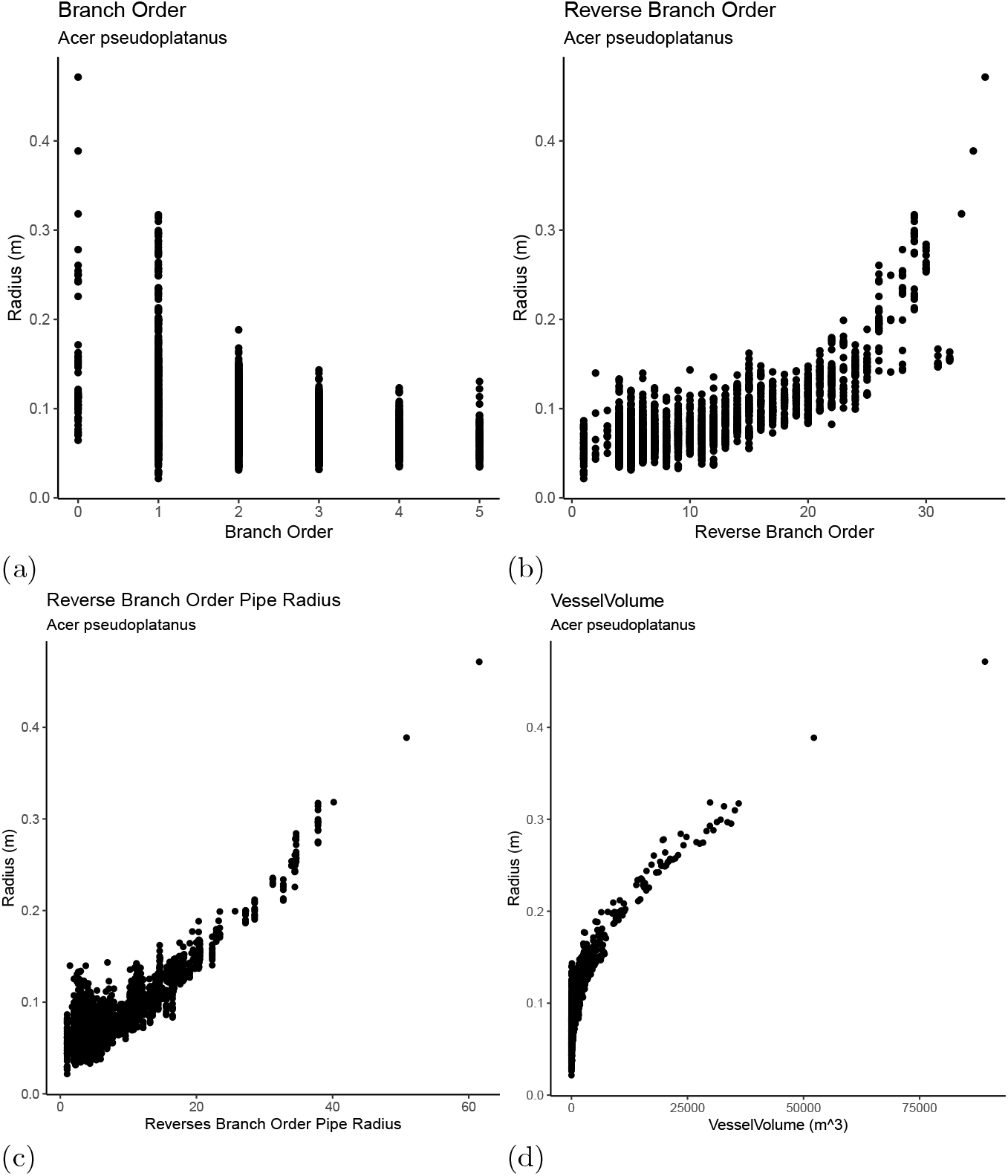
(a) We see the correlation between the traditional `BO` and the `Radius` is not strong. Nevertheless the maximum value per branch order reveals a parabolic pattern. (b) We see the correlation between the `RBO` and the `Radius` is stronger than the one depicted in (a). We see a noisy non linear pattern. (c) When looking at the scatter plot of the `RBOPR` vs the `Radius` we can see a linear pattern, less noisy than the pattern in (b) (d) A power function form non linear pattern is visible when we plot the `VesselVolume` vs the `Radius`. This pattern is the best in the sense that we barely see noise here.

We can see a stronger correlation for the `RBO` compared to the `BO` if we look into the scatter plots of `BO` vs `Radius` and `RBO` vs `Radius`.

There is a proxy potential with a linear pattern in figure 6 (c) on non log transformed input data.

In contrast to the `GrowthVolume` the `VesselVolume` is not affected by over-estimated `Radii` and its correctness relies purely on correct topology. It shows a sharp curved pattern in 6 (d).

### 2.3 QSM Filters

We are about to present two useful filters relying on the `Radius` prediction capabilities of the presented tree measures. No matter if linear or non linear fitting techniques are utilized it is a good behavior to prepare the QSM data. We want to detect cylinders with overestimated `Radii` and not use the measurements of those cylinders in the following fitting routines. As overfitting during the QSM generation is more likely to happen at the tip regions [50, 51], we can simply create one filter to get rid of overestimated cylinders by thresholding the `RBO`. For example Requesting the the `RBO` of valid cylinders has to be larger one removes all tip segments.

The remaining cylinders can then be piped through a statistical outlier filter based on the cylinder’s distance to the aligned TLS sub-cloud.

#### 2.1.3 Linear RBOPR filter

The smaller the `RBOPR` the larger the `Radius` residual is. So we might conclude to not have a homoscedastic, neither a heteroscedastic error. But we know from earlier works that the modeling error for tips is already caused by the input point data itself. Co-registration errors or the foot print size of the Laser have a significant error impact [51]. As the true ground truth tree structure of tip regions is only reflected to some extend in the input clouds we conclude we cannot make an assumption on the underlying distribution of natural `Radius` variation.

We will build the `RBOPR` filter to correct overestimated `Radii` with equation 10.

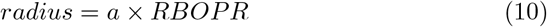

We only correct `Radii` if they differ by more than a user selected percentage deviation threshold from their predicted `Radii`.

#### 2.3.2 Non linear VesselVolume filter

As with the `GrowthVolume` we can try to predict the radius from the `VesselVolume` by a simple non linear power fit as we see in figure 6 (d). So we will use equation 11 to correct cylinders `Radii` and will refer to as the `VesselVolume` filter. To get appropriate initial parameters required for non linear solvers we use the Huxley method on log transformed data [8].

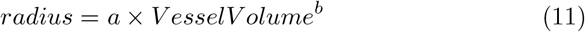

To be in agreement with the geometrical relation between a `Volume` and a `Length` we expect *b* close to the cube root 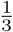.

The solver can still fail, if too large residuals are contained in the fitting data. Non linear fitters are less robust as they can find false, local minima instead of global minima in the error distribution. To get robust against such failiures we recommend to first apply the Filter based on the RBOPR with a large deviation threshold. The linear correction will be applied to the cylinders with the largest errors, but a lot of unmodified cylinders are still usable to predict the `VesselVolume` power function parameters.

The allowed deviation between measured and predicted before we apply a `Radius` correction is user selected in same manner as in the `RBOPR` filter routine.

### 2.4 Data

The author’s of [52] destructively harvested 65 individuals of the species *Fagus sylvatica, Larix decidua, Pinus sylvestris, Fraxinus excelsior*. From `AGB` measurements and density analysis `Volume` reference data was derived. Then the authors produced 65 QSMs with TreeQSM version 2.4.1 [53]. For TreeQSM models a tapering filter [54] relying on a parabolic function to correct overestimated leaves was applied. Both TreeQSMs and cloud data was published at Zenodo [55].

QSMs have been produced as well with SimpleForest v 5.3.1 [44, 56]. We applied the following pipeline on the input clouds:

1. apply a statistical outlier filter on the input clouds [57],
2. produce QSMs with the SphereFollowing method wrapped in an automated parameter search [43, 58],
3. apply the `RBOPR` filter,
4. apply the `VesselVolume` filter.

## 3 Results

Results in this section have been produced with R-Software [59] and for the plots we used ggplot [60].

### 3.1 Comparing against the QSM state of the art

As we have destructively collected reference `Volume` data [55] we can analyze the accuracy of QSM `Volume` predictions. In figure 7 we use SimpleForest QSMs which have been improved by either the linear `RBOPR` filter (a) or the non-linear `VesselVolume` filter (b). We plot as well the results produced with TreeQSM as published [55] (c).

**Fig. 7.**
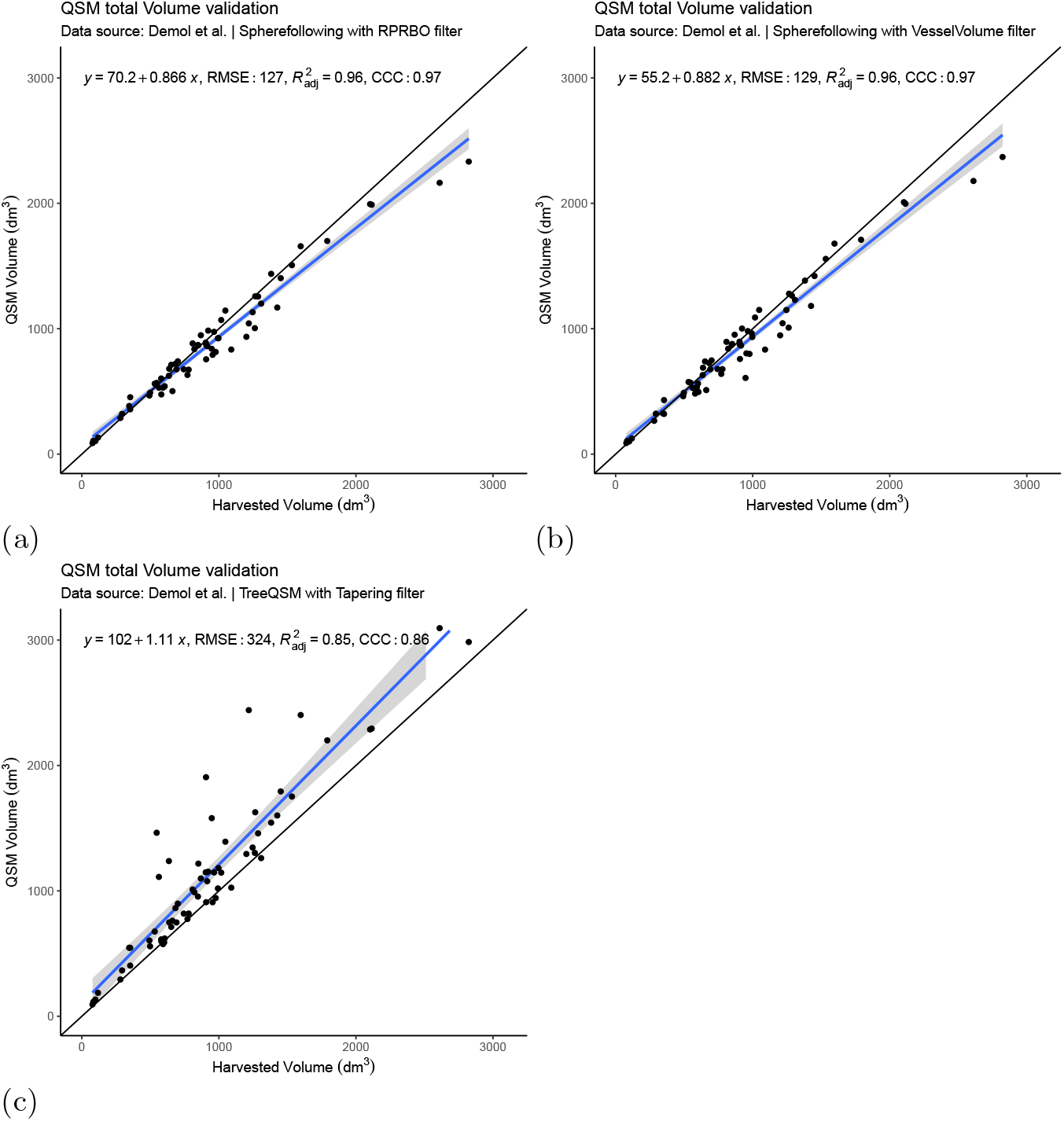
We use free accessible data of tree point clouds, those trees’ destructively collected `Volume` ground truth data and TreeQSM computed `Volume` data [41]. (a) SimpleForest QSMs filtered with the `RBOPR` filter have their `volumes` compared to destructive ground truth measurements. (b) Similar to (a), but this time the `VesselVolume` filter was applied to SimpleForest QSMs. (c) We also show the scatter plot of QSMs stored in the data base. Those have been modeled with TreeQSM algorithm [1] and then filtered by the TreeQSM tapering method [54].

All results are written with their root mean squared error (`RMSE`), their r^2^_adj._ and their Concordance Correlation Coefficient (`CCC`) into table 1.

**Table 1.** Numerical validation of QSM volume measures compared to harvested reference volume measures.

| Software Filter | RMSE($dm^3$) | $r_{adj.}^2$ | $CCC$ |
| --- | --- | --- | --- |
| SimpleForest **VesselVolume** | 129 | 0.96 | 0.97 |
| SimpleForest **RBOPR** | 127 | 0.96 | 0.97 |
| TreeQSM **Tapering** | 324 | 0.85 | 0.86 |

### 3.2 Applying and adapting the Huxley protocol

We took the output of the `VesselVolume` filtered pooled QSM data and looked into the pooled data applying the Huxley protocol. We used as input only data of those cylinders, which did not have been modified by the filter. An unmodified cylinder means that the `Radius` is still the least squares fitted radius. For all trees we still had 13965 cylinders left and those are shown in figure 8. At first glance the untransformed data (a) looks quite diverse. While we can see species clustering also within species the tree individuals show unique patterns. The linear model on the log transformed data (b) shows already a species wise correlation quantified by r^2^_adj._ ranging between 0.89 and 0.96. But if we normalize the `VesselVolume` and the `Radius` to scale for each tree between 0 and 1 a strong pattern is revealed in the not log transformed data (c). The r^2^_adj._s of the species wise linear models range from 0.96 to 0.97 and the slopes of the fitted lines are quite similar in a range of 0.34 to 0.36 (d).

**Fig. 8.**
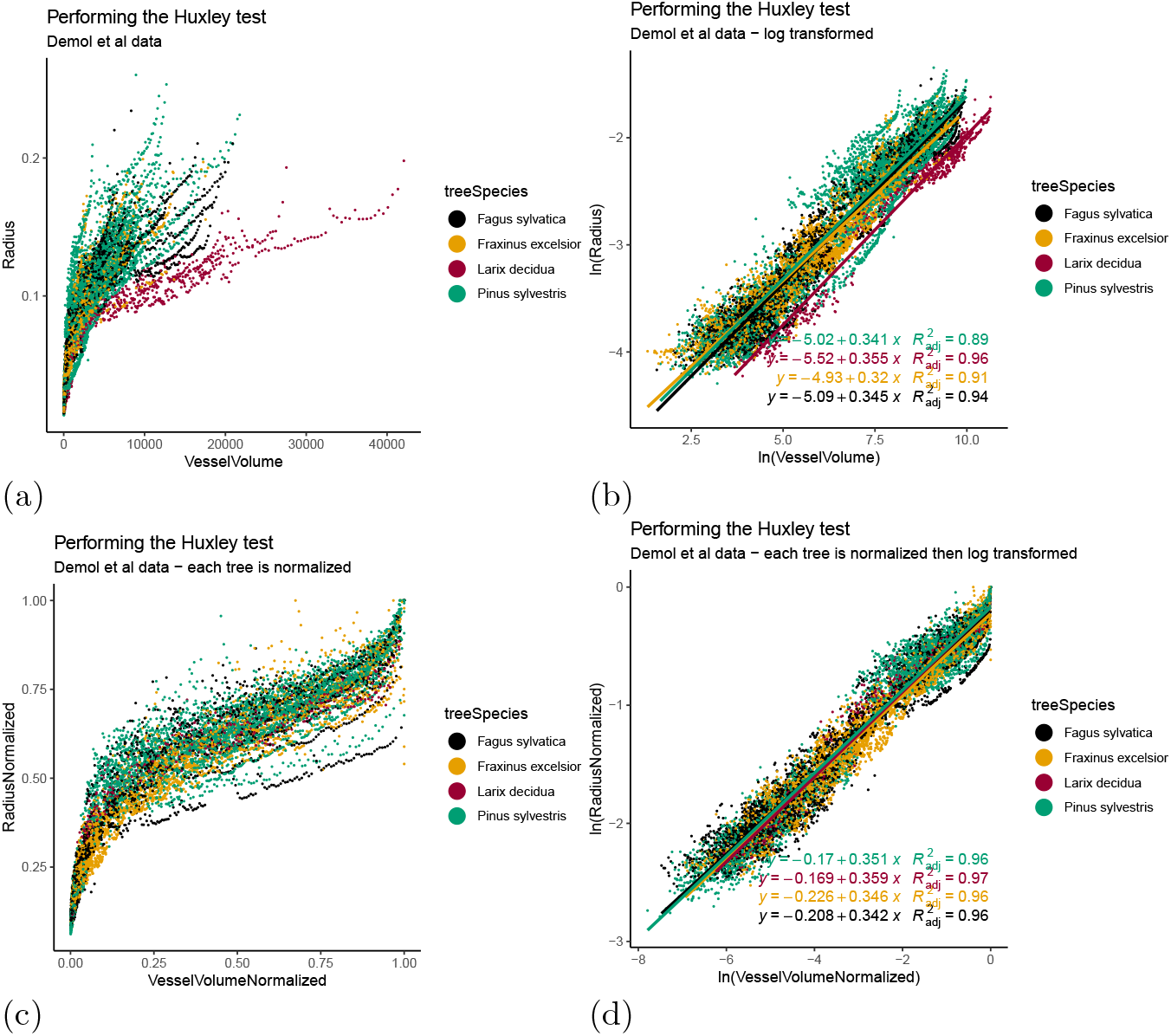
(a) The scatter plot of `Radius` vs `VesselVolume` of 13965 cylinders originated of SimpleForest QSMs produced with open point cloud data [41]. (b) The scatter plot of the log transformed data of (a) with linear models fitted per species. (c) The data of (a), but we normalized each QSM before pooling to have the `VesselVolume` and the `Radius` scale between 0 and 1. (d) After a log transform of data in (c) we fit again linear models per species.

### 3.3 WBE Benchmark

Enquist published values of allometric exponential power coefficients for various tree parameter relations [34]. A few of them we can model with QSM data. Enquist uses as input variable `AGB` which we only approximate with the `GrowthVolume` in this work. We excluded here as well modified cylinders and plotted four allometric relations based evaluated on more than 10’000 measurements in figure 9. We pooled all species together and normalized all parameters to scale between 0 and 1 for each tree.

**Fig. 9.**
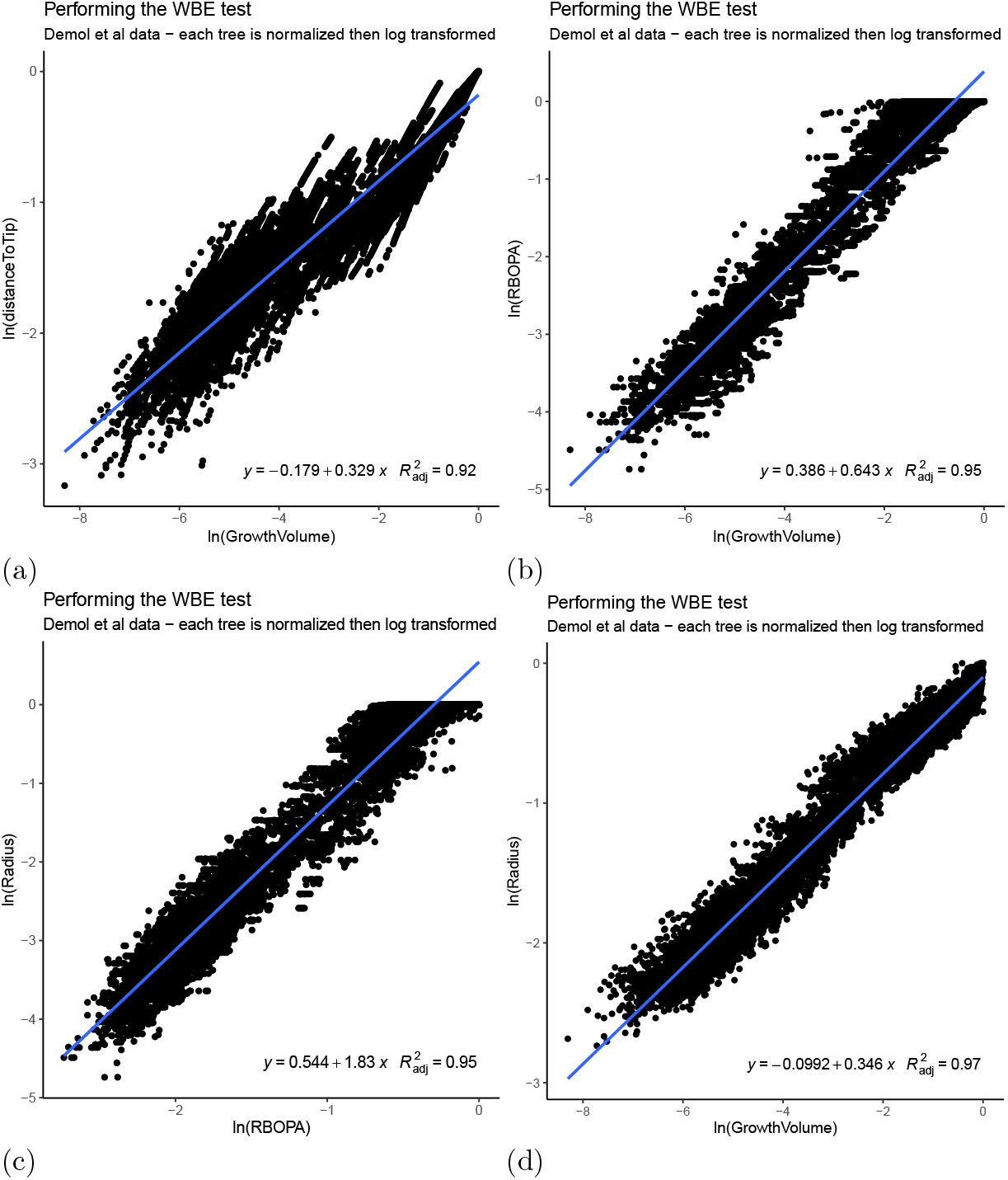
All 65 `QSM`s have been filtered with the `RBOPR` method. We extracted 12352 cylinders which had not been modified by the filter. All of those are cylinders fitted into the point cloud [2] without any further statistical processing. We perform on all data a log transform. (a) The scatter plot of `GrowthVolume` vs `DistanceToTip`. (b) The scatter plot of `GrowthVolume`. (c) The scatter plot of `RBOPA` vs `Radius`. (d) The scatter plot of `GrowthVolume` vs `Radius`.

`PlantMass`, `NumberOfBranches`, `BranchLength` and `Radius` are part of the parameter set for which Enquist predicts the allometric scaling exponents. We compare our estimates with Enquist’s in table 2 where we use:

**Table 2.** We compute for four allometric models the allometric exponent by using the QSMs and compare those exponents to expected published values [34].

| Allometric relation | $r_{adj.}^2$ | $b_{QSM-predicted}$ | $b_{Enquist-predicted}$ |
| --- | --- | --- | --- |
| *DistanceToTip – GrowthVolume* | 0.92 | 0.33 | 0.25 |
| *RBOPA – GrowthVolume* | 0.95 | 0.64 | 0.75 |
| *Radius – RBOPA* | 0.95 | 1.83 | 2 |
| *Radius – GrowthVolume* | 0.97 | 0.35 | 0.38 |

1. `GrowthVolume` as a proxy for `PlantMass`,
2. `RBOPA` as a proxy for `NumberOfBranches`,
3. `DistanceToTip` as a proxy for `BranchLength`.

### 3.4 New allometric functions

In figure 10 we show a four relations which cannot be modeled with parameters measured with traditional methods. We rely in all functions on either the `VesselVolume`, the `GrowthVolume` or the `GrowthLength`.

**Fig. 10.**
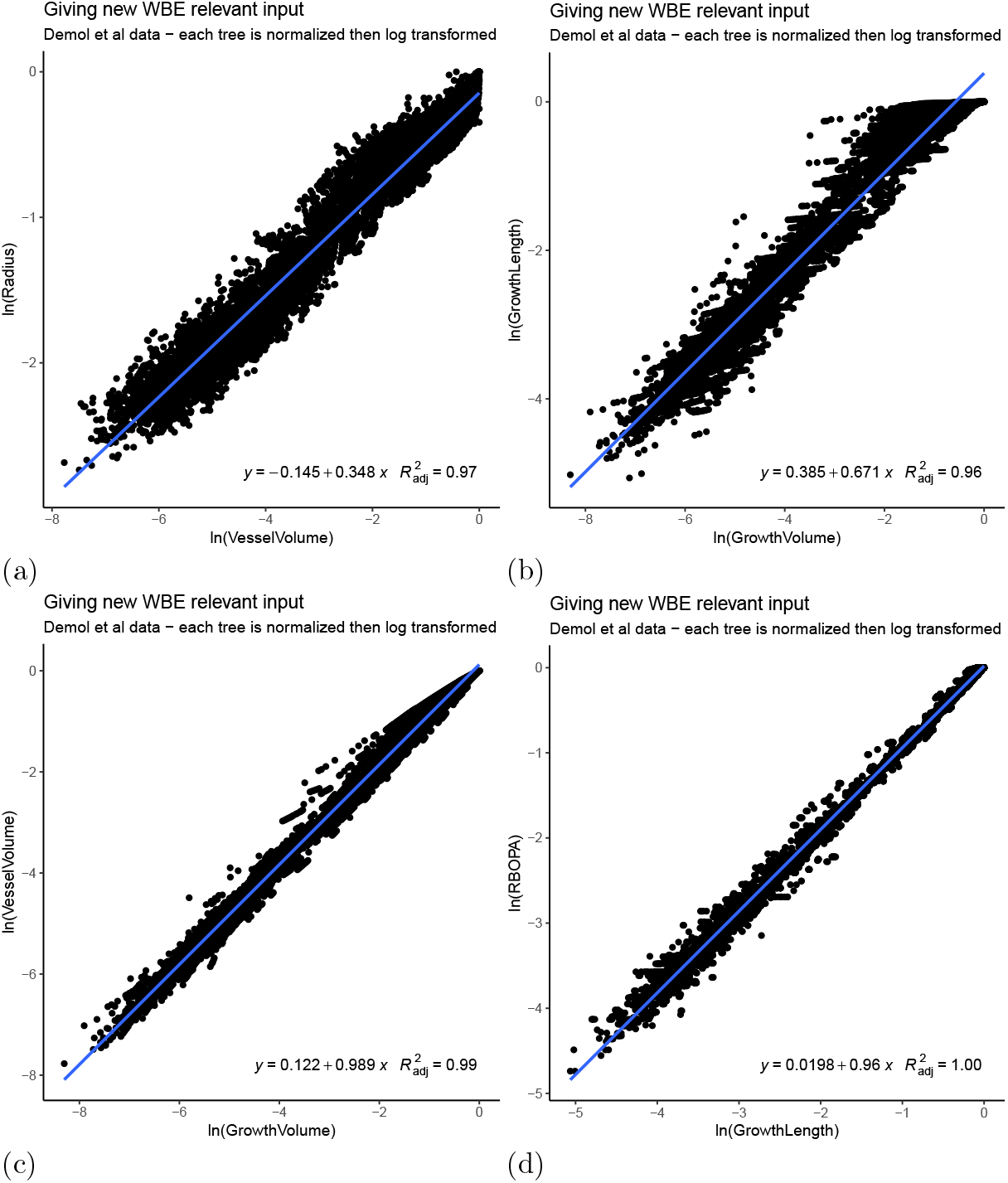
Same data as presented in figure 9. (a) The scatter plot of `VesselVolume` vs `Radius`. (b) The scatter plot of `GrowthVolume` vs `GrowthLength`. (c) The scatter plot of `GrowthVolume` vs `VesselVolume`. (d) The scatter plot of `GrowthLength` vs `RBOPA`.

The results are written in table 3.

**Table 3.** We show four allometric models utilizing our invented parameters and their r^2^_adj._ values.

| Allometric relation | $r^2_{\text{adj.}}$ | $b_{QSM-predicted}$ |
| --- | --- | --- |
| *Radius – VesselVolume* | 0.97 | 0.35 |
| *GrowthLength – GrowthVolume* | 0.96 | 0.67 |
| *VesselVolume – GrowthVolume* | 0.99 | 0.99 |
| *GrowthLength – RBOPA* | 1.00 | 0.96 |

### 3.5 Additional details in un-pooled data

We can see in figure 8 a) and c) sigmoid shapes. More details are revealed in figure 11. It depicts a QSM build from a *Fraxinus excelsior* cloud and a sigmoid function might be more appropriate compared to a simple power function. But as we also see the potential to split the data into two parts and we prefer model each part with an individual power function. The red colored stem cylinder parameters are all cylinders with `BO` zero which start coordinate is lower than the `CrownBaseHeight`. r^2^_adj._ numbers on the log transformed scale are 0.95 for the crown, but only 0.85 for the stem.

**Fig. 11.**
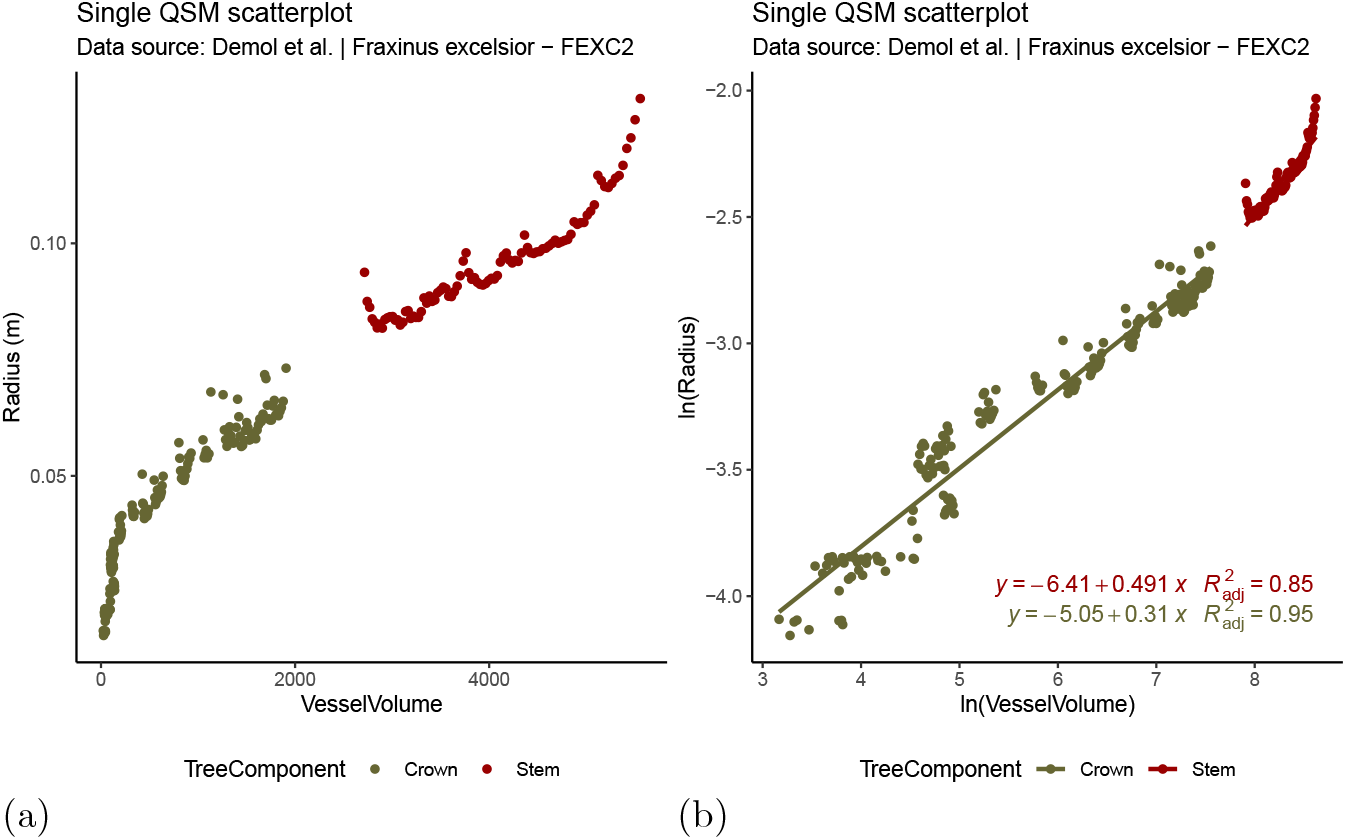
We see cylinder measurements of one QSM of a *Fraxinus excelsior* individual - data base name FEXC2. The QSM was filtered with the `VesselVolume` filter. All modified cylinders have been removed from the analysis. The green dots correspond to branch cylinders and the red to ones to stem cylinders. (a) The whole curve can be described with a s-shaped temporal growth function or split into two data sets and fit two power functions separately for lower stem and crown respectively. Approaches to fit multiple lines into quasi-linear parts have been done already in the early years of allometric research [8]. (b) In fact only the crown seems to follow a linear pattern, the nature of the stem points reveals a curvilinear pattern. The statement is undermined with r^2^_adj._ values of 0.95 for crown, but only 0.85 for fitting a linear model into the stem points.

#### Definition 8

(CrownBaseHeight) The crown starts with the largest branch and we can detect the largest branch inside a QSM by looking for the cylinder which has the largest `GrowthLength` of all cylinders with a `BO` larger zero. The z-coordinate of that cylinder’s start point is the `CrownBaseHeight`.

## 4 Discussion

### 4.1 QSM Volume Accuracy

We imported here in our analysis results [52, 55] which have been produced with TreeQSM. The TreeQSM pipeline which was utilized by Demol et al. uses an algorithm which has been discussed in detail [1]. To achieve an automated parameter search that algorithm was embedded into optimization techniques discussed in [43]. The QSMs then were filtered with the tapering method [54].

Importing the same clouds we also produced SimpleForest QSMs relying on the same parameter optimization technique [43] but utilizing a different algorithm - the spherefollowing algorithm [2]. The output QSMs have then been filtered with the here proposed allometric theory based filters. We fitted linear models into the ground truth `Volume` vs QSM `Volume` predictions, see figure 7 and table 1.

Our results produced data where the `RMSE` is 39 % of the TreeQSM error. The devation of the r^2^_adj._ value from the best score 1.0 is for SimpleForest only 27 % of the TreeQSM derived deviation. For the `CCC` the deviation is only 21 % of the TreeQSM deviation.

To get more insight into the error we want to split up the total error into a measurement error (error_ref_) of the destructively collected reference data, the error produced by the algorithm (error_alg_) which is limited by the potential of the filter to reduce exactly this error (potential_fil_), see equation 12.

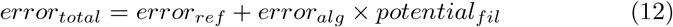

The error_ref_ is the exact same for TreeQSM as well as SimpleForest results. Therefore we will purely focus in the following on error_alg_ and potential_fil_. With the data here used we only show that SimpleForest with one of the allometric scaling filters performs better than TreeQSM with the tapering filters. For splitting up the error further a more complex result matrix would have to be generated.

We conclude that the analyzed tree species can be modeled with the here presented allometric scaling theory based filters. Using our introduced allometric relations performes better than previously published QSM filtering methods [43, 54]. This should help to rise the acceptance to rely on `TLS` scans and `QSM` modeling [61].

### 4.2 Bias

We can see an underestimation of the prediction by SimpleForest, 12 % for `RBOPR` and 13 % for `VesselVolume` filtering. The bias is of roughly the same magnitude like the TreeQSM overestimation (11 %) bias. An untested hypothesis is that in the year of recording less leave growth did happen. Less growing units - the tips which we count - leads to an underestimation of both `RBOPR` and `VesselVolume`.

Time series data recorded under temporal changing weather and climate conditions might open here a complete new field. In hot dry years we might observe underestimation as the sapwood `Radius` and sapwood `Volume` which we use as a predictors are less than in humid and mild years.

### 4.3 Pooling data of different trees

Huxley’s work [8] and follow up publications pool the data of multiple individuals. They use one parameter set per individual. The QSM filters though are based on measurements of one tree only. This is justifiable as QSMs can contain multiple thousand cylinders. Even after our proposed pre-filters to remove over-fitted cylinders enough measurements remain for both the linear and non-linear fit. In fact if we pool all cylinder of the 65 trees we receive more than 12’000 uncorrected cylinders from which we can read a multitude of parameters. All of those cylinders have been fitted with geometrical cylinder fitting routines [2] into the input point cloud data. All cylinders with a statistically modified `Radius` were removed.

We want to test if we can apply the Huxley protocol to pooled data. We sub-divide our measurements per species and plot `Radius` vs `VesselVolume` in figure 8 (a). The plot reveals that individuals of *Larix decidua* have more `VesselVolume` increment per `Radius` increment than individuals of *Fagus sylvatica* have. At first glance the pooling of all trees seems difficult. Even more difficult as we can unambiguously detect individual tree patterns also within one species.

Still we look on the log transformed data (b). In the log transformed domain between and within species variation vanishes a bit. We can fit linear models per species with r^2^_adj._ of 0.89, 0.96, 0.91 and 0.94. Acceptable r^2^_adj._ values but the slope of the models range from 0.34 to 0.36. So the allometric exponents vary a bit and in exponential functions a variation of the exponent has a large impact on the predicted value.

If we normalize our input parameters to range between 0 and 1 by dividing with the maximum value of that parameter per QSM we cannot anymore easily detect tree individual patterns by observation (c).

This statement is undermined by a better r^2^_adj._ ranging from 0.96 to 0.97 and also the slope variation decreased to range between 0.34 and 0.35 (d).

The slope is parameter `b` of equation 11. Our calculated values are in agreement with the previously made assumption that `b` needs to be close to 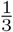. We predict a `Length` from a `Volume` with the cube root.

We conclude that it makes sense to build more complex allometric functions with categorical variables [17], but for the remaining analysis in this work we also found out that normalization is a technique which helps making data of different tree’s or different species better comparable.

### 4.4 How valuable is the WBE model

By comparison with reference data we found out that our filtered QSMs are a representation for the ground truth trees. As not all QSM cylinders are `Radius` filtered and we never modify the topological order in our QSMs by those filters we can collect 12’000 measurements which are derived from our algorithm relying on Gauss Newton cylinder fits [2]. We also found out that by normalization we can pool all those cylinders. We want to use this data to test exemplarily part of the tree modeling theories we reviewed in our introduction.

Some exponents of allometric relations have their allometric parameter published [34]. We compare our empirical predictions against theirs. Using the `GrowthVolume` as a proxy for the `AGB` we can for example build the so important `Diameter` vs `AGB` relation. Enquist predicts here an exponential factor of 0.38 and our predicted value for the `Radius` vs `GrowthVolume` relation is with 0.35 quite close. This is unexpected as our `AGB` proxy ignores all density variation influences and purely relies on the summed up cylinder `Volumes`. The `Radius` vs `RBOPA` relation which to some extend corresponds to the `Diameter` vs `NumberOfBranches` relation deviates only less than ten percent from the Enquist prediction. The `Branchlength` vs `AGB` relation is represented by `DistanceToTip` vs `GrowthVolume` relation. Here the linear model performs worse than the other models. A reason to question this relation on theoretical ground is that we model the relation of a linear path length of the branching structure with a recursive measure which includes side branches. In other words, in that relation predictor and predicted do not represent the same parts of the tree. Our predicted 0.33 is far away from Enquist 0.25. `RBOPA` scales with the `GrowthVolume` with an exponential factor of 0.64. Enquist predicts here 0.75.

### 4.5 Discussing new parameters and their allometric relations

By using scan data of 65 trees we derive 12’000 measurements, ∼100 times more than research relying on traditional data [17, 23]. Also here introduced parameters such as the `RBOPA`, `GrowthLength` or `VesselVolume` can be extracted only from `QSM`s. We show four relations in figure 10 and table 3. The first relation between `Radius` and `VesselVolume` is the one the SimpleTree filter uses. But now we model pooled data of all trees. The r^2^_adj._ is 0.97. Slightly less accurate is the second model for the `GrowthLength` vs `GrowthVolume` relation which has an r^2^_adj._ of 0.96. The performance of scaling between `VesselVolume` and `GrowthVolume` reaches a r^2^_adj._ of 0.99. This relation is already stronger than all the ones we analyzed before. And if we look on the parameters we see that the exponential parameter is close to 1 with 0.99. So we expect to have a linear relation between the area of active pipes and the total area of active and unactive pipes at any point in the tree. `RBOPA` and `GrowthLength` perform similarly well.

### 4.6 Limitations

In figure 11 we reveal that the stem should be excluded from the filters. We assume that a stem should be filtered with a traditional stem taper function and both the `RBOPR` and `VesselVolume` filter should be applied to branches only. Future versions of SimpleForest will provide such functionality.

## 5 Conclusions

### 5.1 Impact on QSM usage

We recommend the usage of one of the two the here presented filtering protocols when working with QSMs. To our knowledge no national forest inventory decided to use QSMs to build `Volume` functions from it up to date. This work provides a significant boost for the `Volume` prediction capabilities of QSMs so we recommend a reevaluation of QSM usage potentials. The computation of linear models is more stable compared to the computation of non linear models. Linear model solvers do not rely on good starting parameters and are always guaranteed to find the global minimum. Therefore we recommend the usage of the `RPOBA` filter over the `VesselVolume` filter.

Generalization of the results should be handled with care. We analyzed here data recorded in leave-off conditions with multiple scans with state of the art equipment and protocols [52]. Only if similar scanning conditions are faced, the accuracy of the results can be expected of the same order as discussed.

Prooning campaigns which are designed to influence a tree’s growth pattern to produce the maximal amount of merchantable wood might result in tree individuals following growth patterns not fulfilling of the here applied Huxley protocol [8]. How well this work performs on either conifers or large evergreen trees needs to be analyzed as well.

### 5.2 Impact on forest ecology theories

The `RBOPA` we interpreted as a proxy for the area of sapwood at any given point in the tree. Of course we could also think of multiplying the `RBOPA` with the expected number of leaves on a new growth unit and derive the total number of leaves. So the `RBOPA` can be seen as a proxy for the area leave index as well.

With open QSM data bases tree modelers have from now on a much more convenient way to collect data.

### 5.3 Future Work

We first want rework the preprinted SimpleForest software paper to provide an up to date reference for the here used free software tool [44].

We also identified the need for a data paper about an open data-base-system of. Various data sets have already been made accessible [50, 52, 62] and we want to combine all small data bases to a larger one backuped by a manuscript. We hereby would like to convince more data owners to share their data. Instead of 65 trees future analysis can be run on few hundred of tree clouds with harvested volume data.

We like to envisage a collaborative paper with TreeQSM and 3dForest personnel. As soon as the work on the data base paper is done we could run all three softwares unfiltered and combine each result with one existing filter technique.

## acknowledgements

We would like to acknowledge Matthias Disney for providing the SimpleForest development with constructive feedback over the last decade via emails and manuscript reviews.

## Declarations

### Funding

All development of SimpleForest software including the presented research results have been private funded by Jan Hackenberg. The authors declare that no funds, grants, or other support were received during the preparation of this manuscript.

### Conflict of interest/Competing interests

The authors declare no conflict of interest.

### Availability of data and materials

The `TLS` clouds and the TreeQSM results can be downloaded [55].

### Code availability

We use for the computation in addition to the mentioned data the persistent SimpleForest release v5.3.1 [56]. Here a precompiled windows executable exists.

The code of SimpleForest can also be downloaded from the Git repository https://gitlab.com/SimpleForest/computree/.

SimpleForest QSMs as well the R scripts for the statistical analysis and plots are embedded in the manuscript repository https://gitlab.com/SimpleForest/allometricscalingpaper.

### Authors’ contributions

Jan Hackenberg developed the SimpleForest software as well as the methods and algorithms. He did the statistical analysis and the writing of the first version of the manuscript. Jean-Daniel Bontemps helped improving the allometric methods of SimpleForest and did the review of this manuscript.

## Notes

https://zenodo.org/record/5131717/

### Competing Interest Statement

The authors have declared no competing interest.

### Summary of Updates

We improved the section about recurisve topology information.

